# Assembly of Protein Complexes In and On the Membrane with Predicted Spatial Arrangement Constraints

**DOI:** 10.1101/2023.10.20.563303

**Authors:** Charles Christoffer, Kannan Harini, Gupta Archit, Daisuke Kihara

## Abstract

Membrane proteins play crucial roles in various cellular processes, and their interactions with other proteins in and on the membrane are essential for their proper functioning. While an increasing number of structures of more membrane proteins are being determined, the available structure data is still sparse. To gain insights into the mechanisms of membrane protein complexes, computational docking methods are necessary due to the challenge of experimental determination. Here, we introduce Mem-LZerD, a rigid-body membrane docking algorithm designed to take advantage of modern membrane modeling and protein docking techniques to facilitate the docking of membrane protein complexes. Mem-LZerD is based on the LZerD protein docking algorithm, which has been constantly among the top servers in many rounds of CAPRI protein docking assessment. By employing a combination of geometric hashing, newly constrained by the predicted membrane height and tilt angle, and model scoring accounting for the energy of membrane insertion, we demonstrate the capability of Mem-LZerD to model diverse membrane protein-protein complexes. Mem-LZerD successfully performed unbound docking on 13 of 21 (61.9%) transmembrane complexes in an established benchmark, more than shown by previous approaches. It was additionally tested on new datasets of 44 transmembrane complexes and 92 peripheral membrane protein complexes, of which it successfully modeled 35 (79.5%) and 15 (16.3%) complexes respectively. When non-blind orientations of peripheral targets were included, the number of successes increased to 54 (58.7%). We further demonstrate that Mem-LZerD produces complex models which are suitable for molecular dynamics simulation. Mem-LZerD is made available at https://lzerd.kiharalab.org.

## INTRODUCTION

Protein-protein interactions are fundamental to many biological processes in living cells. Membrane proteins play an essential role in many of these processes, where they act as gateways of cellular signaling pathways, pumps, and more, facilitating selective transport processes across membranes. To understand the detailed mechanisms of these processes, modeling the 3D structures of their associated protein complexes is a critical step. While protein complex structures are steadily being determined by experiment and deposited in the Protein Data Bank (PDB) [1, 2], experiments are still expensive and a substantial human and instrument time investment, assuming the proteins at hand are compatible with the methods available [3]. Moreover, structures of protein complexes are often extremely difficult to determine by experiments, even without the involvement of a membrane. Thus, when a protein complex structure has not yet been experimentally determined, computational tools can be used to construct atomic models [4]. A protein docking program can take component proteins, called subunits, as input and assemble them into models of the protein complex. Many general protein docking methods and specialized versions thereof have been publicly released, such as ZDOCK [5], HADDOCK [6], ClusPro [7], RosettaDock [8], HEX [9], SwarmDock [10], and ATTRACT [11]. Even protein structure prediction methods like AlphaFold [12] have been tweaked to be able to output multimeric structures [13]. The rigid-body docking method LZerD [14–17] in particular has been consistently ranked highly in the server category in CAPRI [18, 19], the blind communitywide assessment of protein docking methods.

The fact that proteins in vivo are not generally interacting in isolation in a uniform environment is often confounding to computational modeling of complexes. Even with state-of-the-art modeling techniques, existing docking methods struggle to rigorously handle environments other than a uniform aqueous environment [20, 21]. Membranes create an environment where hydrophobic surfaces are not strongly energetically incentivized to be buried in the protein-protein interface. For example, in the transmembrane halorhodopsin protein family as found in halobacteria, which is included in the benchmark of Mem-LZerD, the protein-protein interface is not especially rich in hydrophobic amino acids relative to the remainder of the molecular surface, much of which participates in the protein-lipid interface [22]. Highlighting the utility of docking methods, a halorhodopsin structure for *Halobacterium salinarium* was available in 2000, but no structure for *Natronomonas pharaonic* was available until 2009 [23]. Homology-based methods can be used to model the subunits, and a docking method can then be used to explore the space of interaction poses. Other proteins do not wholly embed in the membrane, but instead pass only partway through or interact only with the membrane surface. These peripheral membrane proteins are likewise important and are implicated for example in sensitivity to membrane composition [24].These broad categories of membrane protein break down further into classes with substantial mechanistic differences, from the purely α-helical transmembrane regions most commonly considered by computational methods, transmembrane β-barrels, to peripheral membrane proteins attaching to the membrane with amphipathic helices [25] or with hydrophobic loops [26]. Transience in certain interactions between these proteins can render them difficult to directly consider in vitro [27], but more accessible via computational modeling [28]. It is then clear that techniques capable of modeling interactions involving membranes have the potential to elucidate many cellular processes in many biological contexts. Detailed mechanistic understanding of membrane protein complexes currently represents a major knowledge gap in molecular biology and is the subject of much active investigation [29]. Recent studies have shown that protein structure prediction methods can enable combinatorial modeling of putative interactions among, for example, the cytochrome c maturation system I proteins of *E. coli* [28].

Several computational techniques have been developed which predict the modes of interaction of proteins in a membrane environment. MPDock, part of the RosettaMP software collection, takes a specification of membrane chemistry and dimensions to model the assembly of transmembrane complexes from bound experimental structures [30]. In their benchmark, the bound subunits were pulled apart, repacked according to the algorithm of Rosetta, and then pulled back together. Memdock carries out rigid-body docking of α-helical membrane proteins by constraining their orientation, before a finer-grained refinement procedure, and makes the assumption that the whole-input center of mass roughly coincides with the membrane midplane [31, 32]. JabberDock for membrane proteins represents the subunits with a volume map incorporating dynamics information from an expensive 70 nanosecond molecular dynamics simulation and uses particle swarm optimization techniques to explore the pose space, which required strict constraints on the pose space [33]. In summary, MPDock requires bound native structures as input and requires manual processing, Memdock was only tested on α-helical proteins and requires assumptions about the membrane location, and is further unable to include soluble regions of proteins in the modeling, and JabberDock requires substantial molecular dynamics calculations and is limited to exploring a tight region of the search space the precludes tolerance for misorientation. These existing analyses require orientations taken from bound complex structures, rather than orientations predicted separately for each subunit. In the context of blind modeling, however, the precise orientations which individual subunits take on upon binding are not known. A method for blind membrane docking should predict the membrane orientations of the input subunits and have a docking search space which is narrow enough to exclude prediction errors, but broad enough to tolerate reasonable errors in orientation predictions.

Mem-LZerD, which we developed in this work, is based on the LZerD rigid-body docking method and its extensions [14–17, 34–36] LZerD uses a soft surface representation of the protein subunits based on geometric hashing [37] and 3D Zernike descriptors (3DZDs) [38–43], which allows for fast generation of docking poses without considering side-chain repacking. The geometric hashing procedure originally used internally by regular LZerD also admits more site-specific orientation and translation restrictions. Mem-LZerD targets any transmembrane protein complexes, as well as peripheral membrane proteins. The geometric hashing data structure used by Mem-LZerD is newly augmented with the positioning of each sample point relative to the membrane generated by the Positioning of Proteins in Membranes (PPM) algorithm [44, 45], as well as the angular orientation in the same membrane. Pruning the search space in this way, skipping infeasible poses, yielded a running time 74 times faster on average compared to regular LZerD without search space or model constraints. This speedup highlights that for membrane proteins, most of the calculation time in regular LZerD is spent on infeasible poses, e.g. those which are sideways or entirely outside the membrane, which then contaminates the output model ranking. We will directly show that the examined regime is inaccessible to ordinary rigid-body docking and that interactions of both transmembrane and peripheral membrane proteins are accessible in this framework. Mem-LZerD yielded acceptable models in the CAPRI criteria within the top 10 models for 13 of 21 (61.9%) unbound docking targets of the Memdock benchmark set, which is a greater fraction than successfully modeled by existing methods Memdock or JabberDock. Memdock itself was able to model only 4 (19%) targets, or 36.4% of the 11 targets for which the protocol modeled unbound subunit structures at all. On that smaller set, Memdock successfully modeled 5 (45.5%) of the targets. Previous studies assumed knowledge of the ground truth subunit orientations, rather than predicting them as in the Mem-LZerD protocol. When assuming the ground truth orientations, Mem-LZerD successfully modeled 14 of 21 targets (66.7%). On our separate transmembrane protein benchmark set, Mem-LZerD successfully modeled 35 of the 44 (79.5%) unbound benchmark docking targets, while on a preoriented peripheral membrane protein benchmark set, Mem-LZerD successfully modeled 54 of 92 (58.7%). Mem-LZerD has been incorporated into the LZerD webserver, available at https://lzerd.kiharalab.org. We further show that the protocol pipeline of the Orientations of Proteins in Membranes/Positioning of Proteins in Membranes (OPM/PPM) suite [44], Mem-LZerD, and CHARMM-GUI [46] can produce ready-for-simulation files of sampled binding poses in explicit lipid membranes.

## RESULTS AND DISCUSSION

### Overview of Mem-LZerD

The docking procedure of Mem-LZerD follows the observation that presence of a lipid membrane restricts the poses two proteins may take on when they assemble into a complex. Mem-LZerD builds assembled complex models by independently orienting input proteins using PPM [44], then applying a geometric hashing-based procedure adapted from LZerD with augmentations that allow pruning the search space to sample membrane-bound poses, before a final clustering and scoring procedure yield the final outputs The overall procedure of Mem-LZerD is diagrammed in Figure 1a.

**Figure 1.**
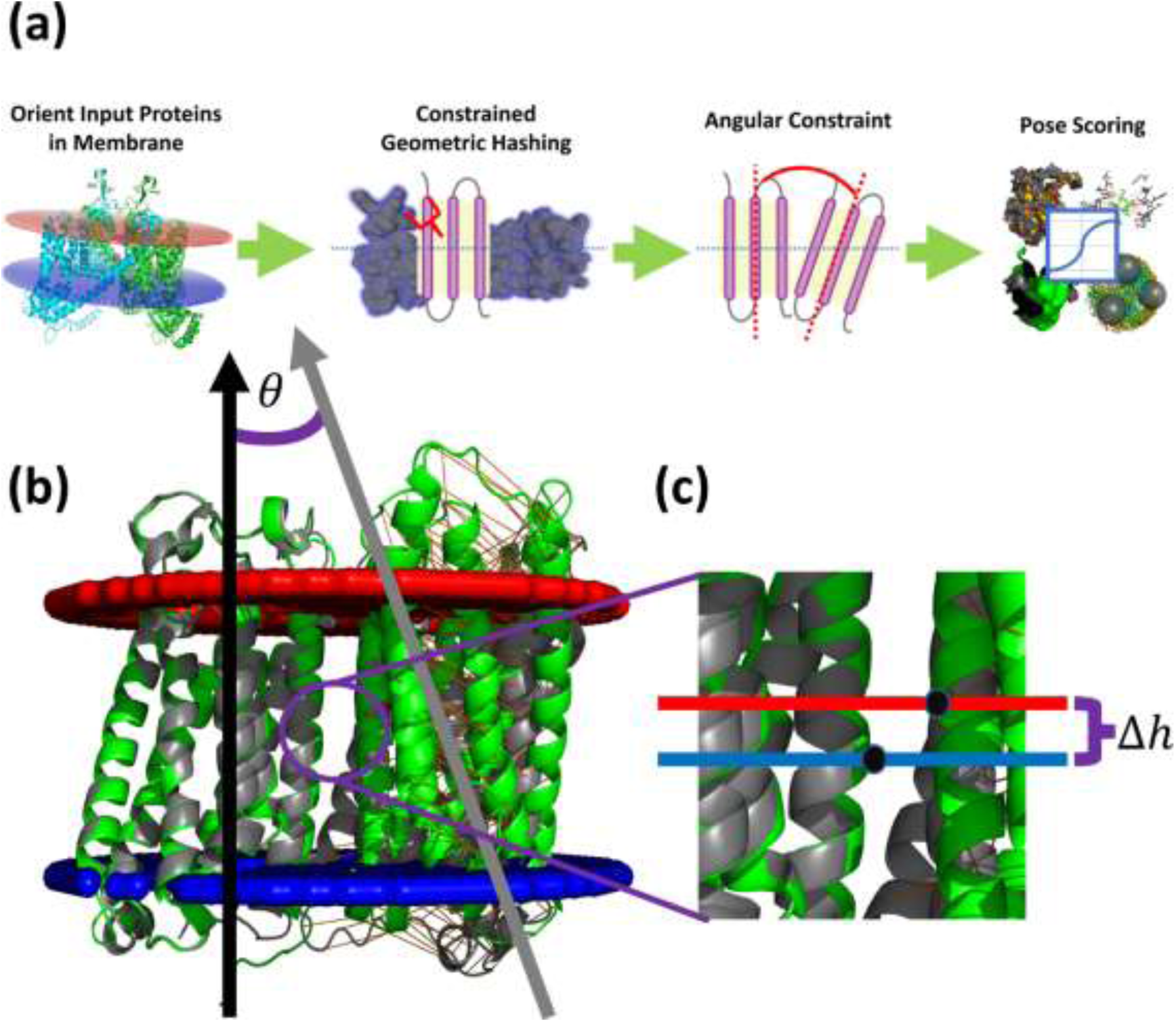
(a) Overall flow of the Mem-LZerD method. **(b)** Tilt constraint used by Mem-LZerD. The black and grey arrows represent example the axes perpendicular to the membrane planes as oriented by PPM, while 𝜃 is the tilt angle between those axes. The red and blue boundaries represent the inner and outer membrane surfaces predicted by PPM for the receptor subunit. **(c)** The height constraint used by Mem-LZerD. The black dots represent points on the molecular surface sampled by the initial stages of LZerD, while the difference between the pre-docking oriented heights relative to the membrane Δℎ is the height used to prune the docking search.

The first step is to orient both the receptor and ligand protein subunits separately in the membrane using PPM, which minimizes its computed free energy of transfer between the aqueous and membrane environments. PPM has been tested on thousands of membrane proteins and validated against hundreds of experimental results [44]. PPM, however, does not in general generate precisely the orientations the subunits take on in their bound states, but the search and sampling procedure of Mem-LZerD is sufficiently tolerant that such precision is not necessary. Transmembrane proteins are immersed in the membrane and therefore more clearly restrained by it, while peripheral membrane proteins merely interact with the membrane surface and are less clearly restrained.

In the next stage, Mem-LZerD searches the pose space using the LZerD algorithm while accounting for the restricted pose space using an augmented data structure. Unlike the data structures used in original LZerD, the newly augmented geometric hashing in Mem-LZerD facilitates restricting the search space without the expense of running the full pipeline and post-filtering the output. The two subunits are assumed to be oriented to the same membrane. The residuals of the protein surface sample point height, a novel development, and the angles between the transformed membrane planes, as illustrated in Figure 1b, established in the literature and validated in this work, together then constrain the pose space. These simple criteria facilitate efficient search of the pose space without wasting time calculating intermediate scores for infeasible poses.

In Figure 2a and Figure 2b, the distributions in terms of height and tilt of docked models of at least acceptable CAPRI quality for cytochrome c oxidase subunit 1 and subunit 3 (PDB 1M56 chains A and C) are shown for docking with predicted orientations and docking with ground truth orientations, respectively. Both panels clearly illustrate that models of at least acceptable CAPRI quality tend to be found closer to the origin, i.e. docking conformations with a small height difference and tilt angle difference, regardless of whether the basis of the constraints used is a blind or non-blind orientation. To select optimal cutoff values for the height and tilt angle differences, an exhaustive parameter sweep was carried out for height difference cutoffs of 4 Å, 8 Å, and 12 Å and tilt difference cutoffs of 0.2 radians, 0.4 radians, and 0.8 radians on the training set of 20 nonredundant transmembrane protein complexes taken from the OPM database [45] as described in Materials and Methods. For each combination of parameters, the enrichment factor (EF) and recall were calculated where applicable across the targets between the baseline LZerD sampling and the Mem-LZerD sampling with constraints applied. As shown in Figure 2c, a combination of 8 Å height and 0.4 radian tilt differences exhibits the highest EF value before a dramatic loss of recall is observed: these settings resulted in an EF of 20.3 and a recall of 0.27, while the next-highest EF of 33.1 dropped the recall to 0.06. The highest EF of 135.7 would have yielded a recall of only 0.18. These cutoffs also agree with those found in existing literature, although the height referred to in such past work has been the subunit centroid height and not a property of the molecular surface [32]. Thus, while the formulation of height in Mem-LZerD differs substantially from that in existing methods, this parallel illustrates that Mem-LZerD heights can generally be interpreted using the same understanding as of centroid heights. Supplemental Video S1 illustrates the difference between the decoy poses sampled by this procedure and the original LZerD search for an intramembrane aspartate protease complex (PDB 4HYG), using the final chosen constraint cutoffs of 8 Å and 0.4 radians. Using this formulation of constraints, many clearly impossible models, including many sideways or totally dis-immersed poses, which are sampled by LZerD are not sampled by Mem-LZerD. In this way, both the running time of the sampling and the number of raw unclustered decoys were reduced by a factor of 74 on average across our benchmark set. In the case of the decoy distributions shown for target 1M56 in Figure 2ab, the numbers of models were reduced by factors of 66 and 69 for blind and preoriented docking, respectively.

**Figure 2.**
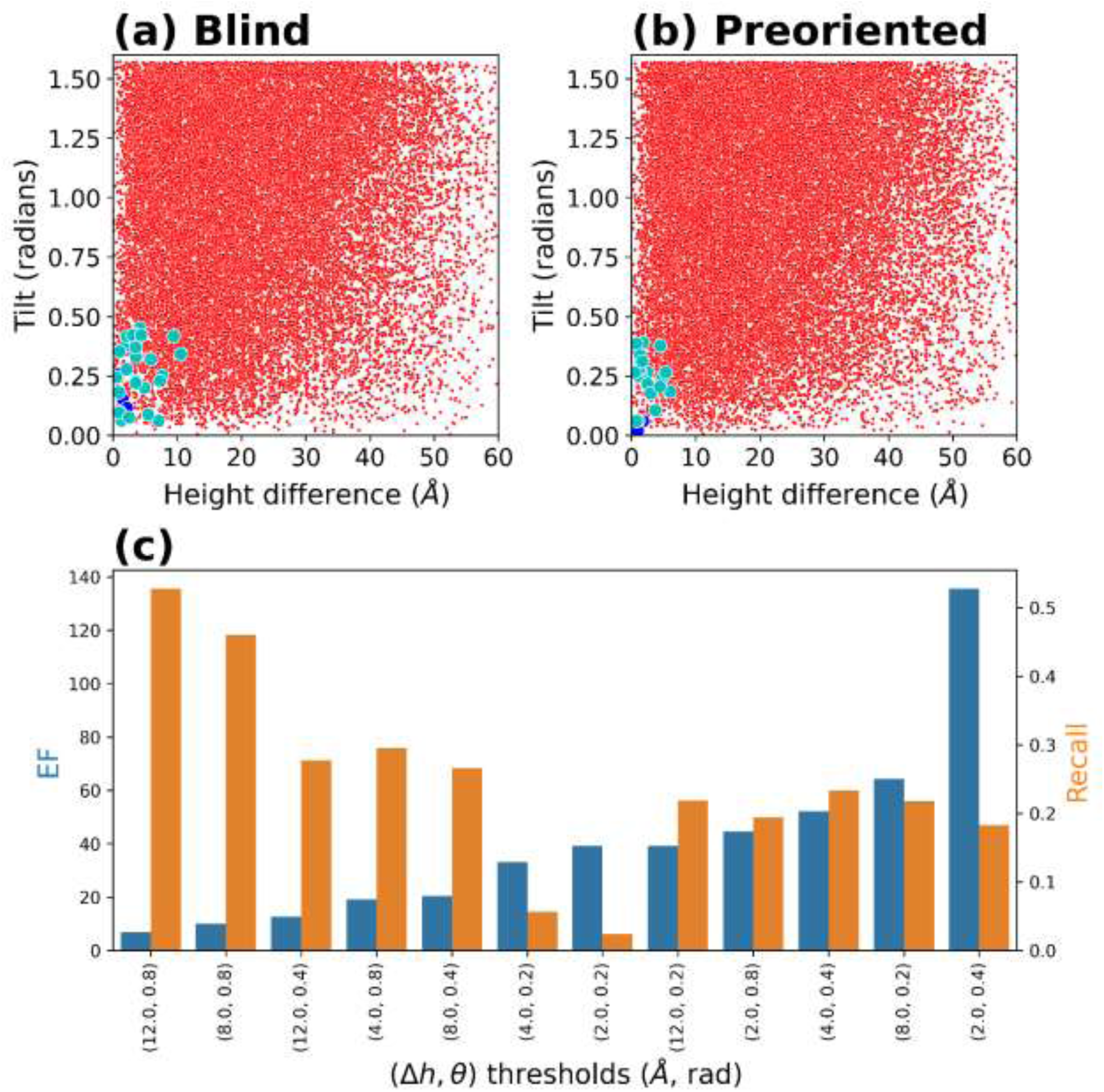
Deviation from ideal membrane spatial positioning and orientation using (a) blindly oriented and (b) preoriented docking inputs for PDB 1M56, as included in the usual top 50,000 models from LZerD. Red: incorrect quality models. Cyan: acceptable quality models. Blue: medium models. Green: high quality models. Even when blindly predicted inputs are used, 0.4 radians tilt and 8 Å height difference are sufficient to capture most models of acceptable or better quality. (c) Validation of the 0.4 radians tilt and 8 Å height difference cutoffs. Blue: enrichment factor of the constrained pose space. Orange: recall of the constrained pose space. When parameter combinations are sorted by enrichment factor, recall sharply declines after 0.4 radians tilt and 8 Å height difference.

### Docking results on the Memdock bound and unbound dataset

Mem-LZerD was trained on a nonredundant set of 20 transmembrane protein complexes and benchmarked on nonredundant sets of 21 and 44 transmembrane protein complexes originally used by Memdock [32] and newly constructed for Mem-LZerD, respectively, as described in Methods. Mem-LZerD was additionally benchmarked on a nonredundant set of 92 peripheral membrane protein complexes. The main evaluation results are summarized in Table 1 and Table 2, covering the existing benchmark set [32] and our new benchmark set, respectively. Modeling for a target was considered successful if one of more of the top 10 output models was of at least acceptable CAPRI quality. According to the standard CAPRI criteria, a model is acceptable if it has a fraction of native contacts (𝑓_nat_) of at least 0.1 and either an interface RMSD (I-RMSD) of at most 4.0 Å or an L-RMSD of at most 10.0 Å [18, 21, 47, 48]. A rank-of-first-hit (RFH) value (as shown in Table 1) that is as high as 10 is considered a success for the given method on the given target.

**Table 1.**
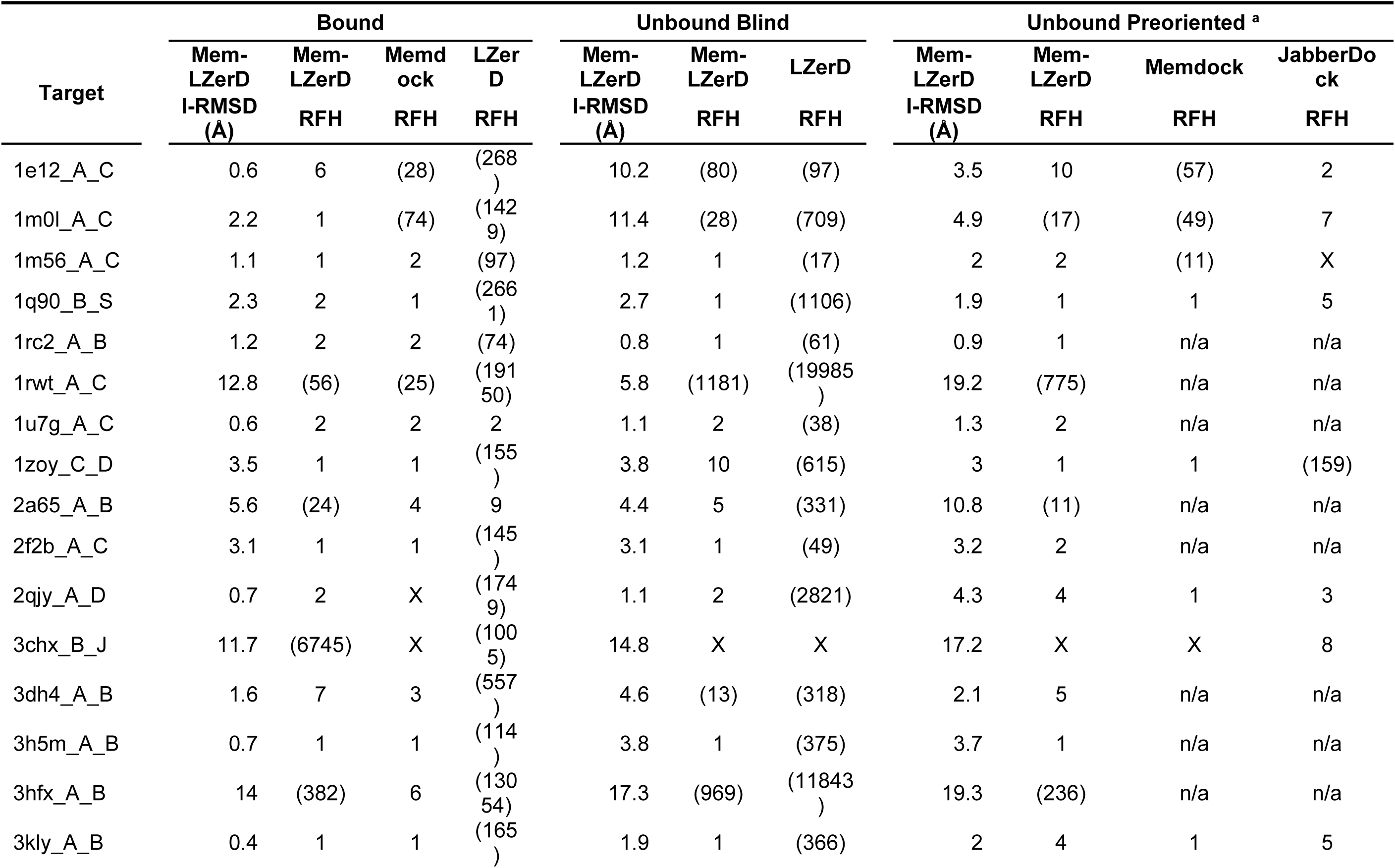
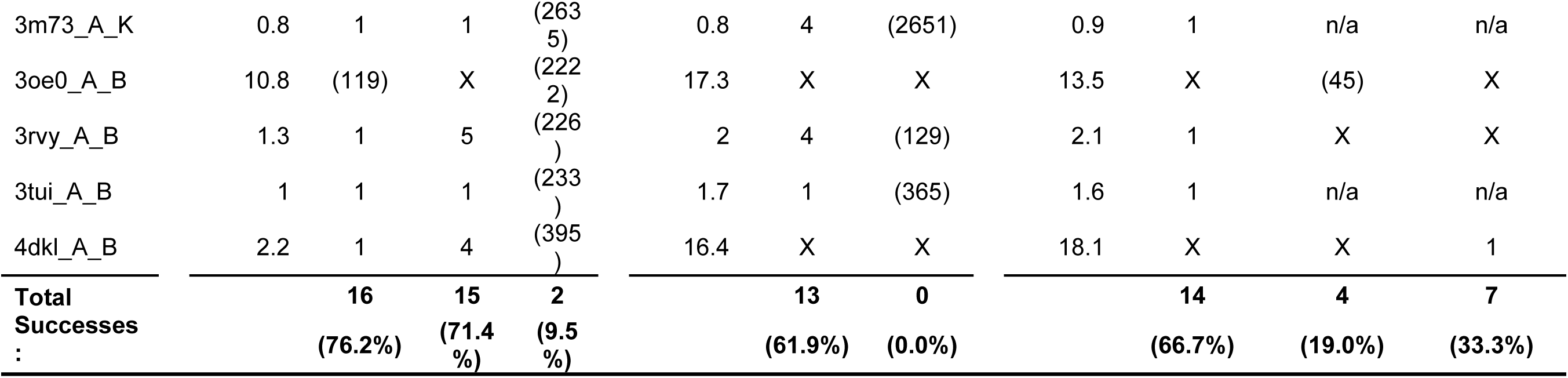
Docking performance of the 21 individual targets of the Memdock benchmark set. RFH is the rank of the first CAPRI-acceptable model generated, if any, for a given target. RFH must exist and be at most 10 for modeling to be considered successful. n/a indicates that there was no unbound model modeled for a target in the published study. X indicates that there no docked models of at least acceptable quality in the analyzed output; for Mem-LZerD, this is limited to 50,000, the pre-clustering model cutoff; for Memdock, this is limited to the top 100 models. We considered only the top 10 models from either method in our evaluation, but scored model ranks worse than 10 are shown here for completeness in parentheses. The I-RMSD columns list the best interface RMSD among the top 10 models. Separate preoriented docking outcomes are not show for regular LZerD runs since the original LZerD pipeline does not consider input model orientation. **^a^** These evaluations of Memdock and JabberDock from the paper first describing their use for membrane docking used premade orientations from OPM, which are orientations of the full bound complexes rather than the subunits independently. The Mem-LZerD unbound benchmark otherwise shown is fully unbound docking, with membrane orientations calculated by PPM separately for each subunit without regard to the full complex. However, here we additionally show Mem-LZerD results when using ground truth orientations as in the original Memdock and JabberDock studies.

**Table 2.**
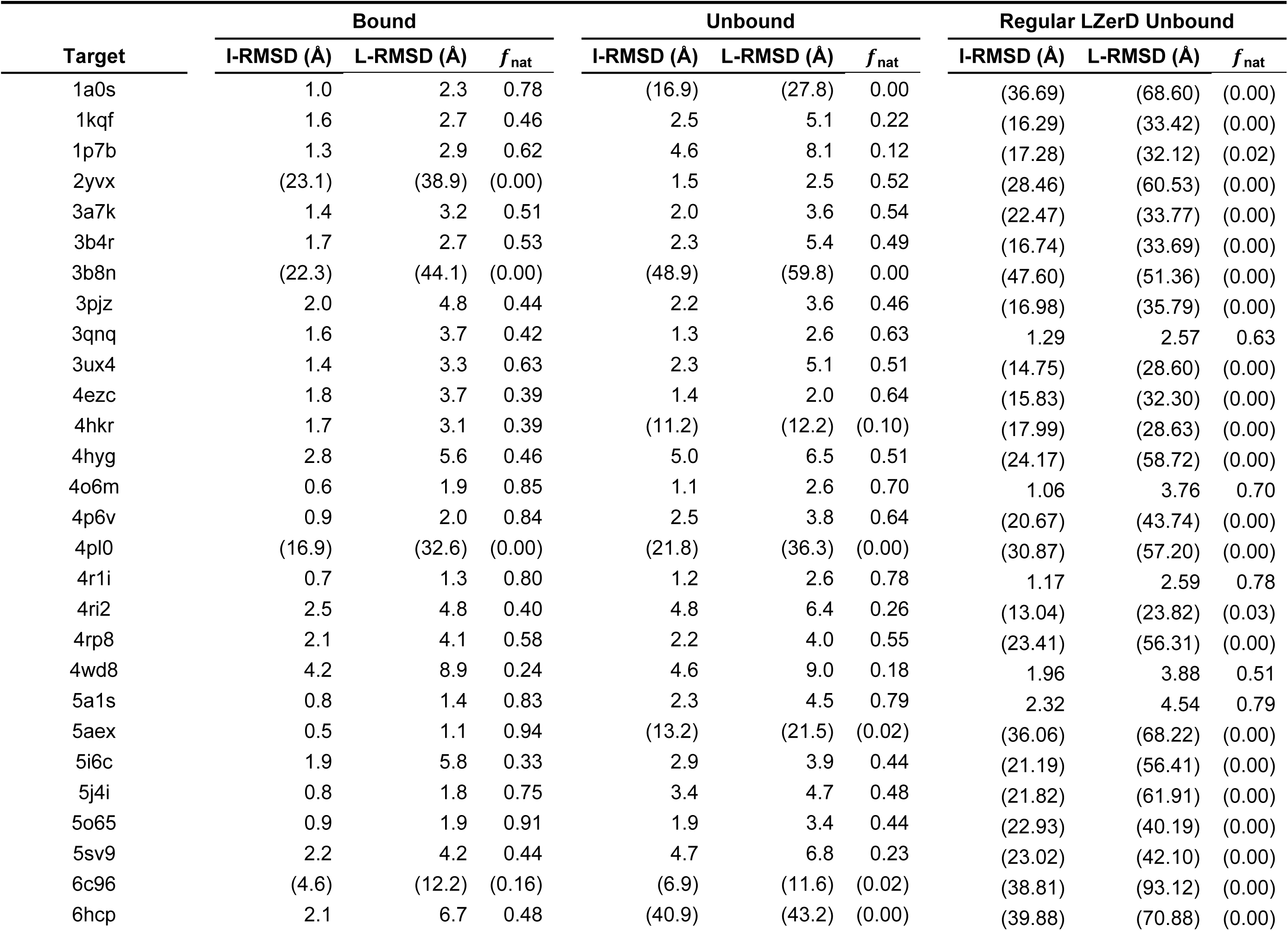
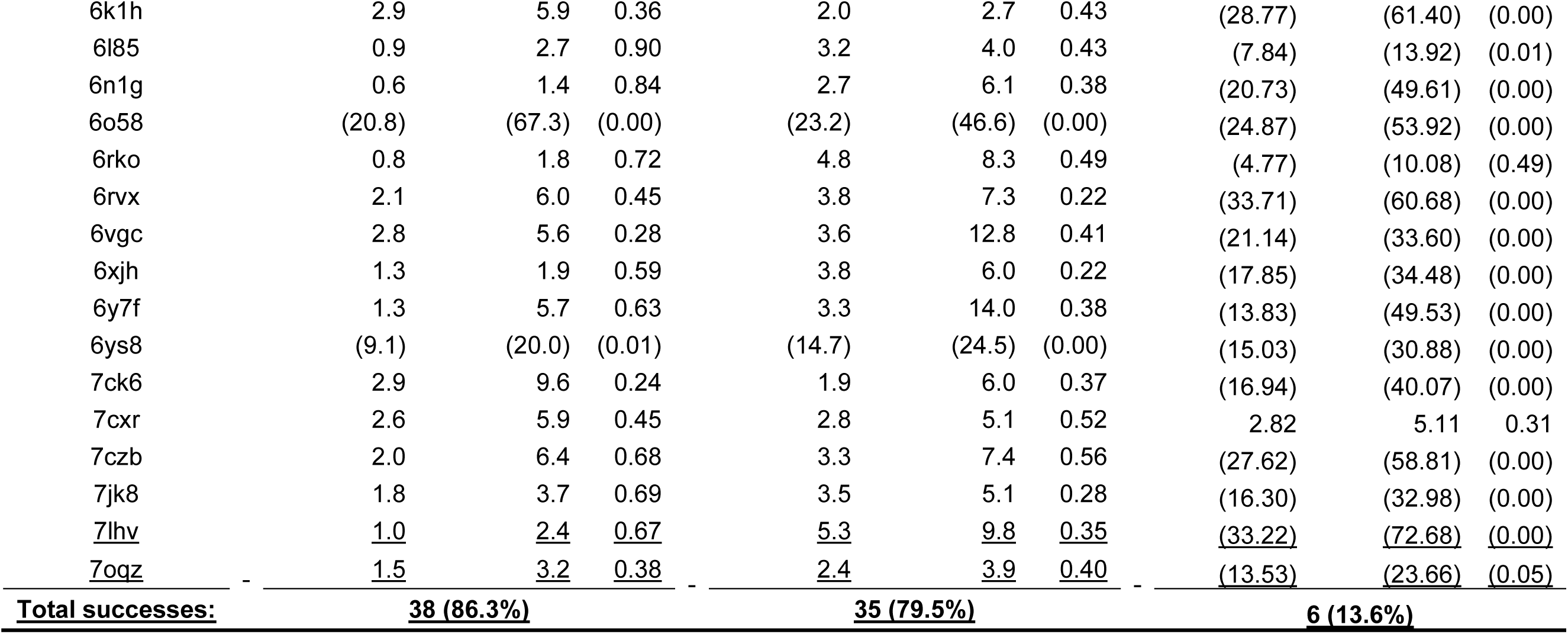
Docking performance of the 44 individual targets of the Mem-LZerD transmembrane protein benchmark test set. Parentheses indicate that no models of at least CAPRI-acceptable quality were ranked within the top 10. The columns list the best of each measure among the top 10 models.

To examine the precision of docking using Mem-LZerD versus past rigid-body methods, we compared the performance of Mem-LZerD against the Memdock benchmark set [32], as shown in Table 1. The Memdock algorithm is designed specifically for α-helical transmembrane proteins, rather than general membrane proteins which may be β-barrels or not transmembrane at all. Thus, we compare using Memdock’s original benchmark set of 21 α-helical transmembrane complexes. Memdock achieved a success rate of 71.4% (15 out of 21 cases) for bound docking, while Mem-LZerD achieved a success rate of 76.2% (16 out of 21 cases; Table 1, Bound column group). On this same benchmark, the baseline LzerD pipeline, which does not consider membrane constraints, succeeded on no targets for unbound docking and 2 of 21 (9.5%) targets for bound docking. Transmembrane docking targets were thus shown to largely require specialized methods, and Mem-LZerD achieved a higher success rate than such existing specialized methods.

For unbound docking on the Memdock benchmark (Table 1, Unbound Blind and Unbound Preoriented column groups), results from past docking experiments using both Memdock and JabberDock were available, and we evaluated the unbound docking performance of Mem-LZerD against these. Here, Mem-LZerD achieved an overall success rate of 61.9% (Table 1, Unbound Blind column group), while Memdock and JabberDock achieved 19.0% and 33.3% respectively overall (Table 1, Unbound Preoriented column group). The Unbound Preoriented column group, compares unbound docking performance when knowledge of the ground truth subunit orientations is assumed. Here, we calculated preoriented unbound input models by superimposing them individually to their native complex structures as oriented in OPM. The preoriented Mem-LZerD success rate was then 66.7% (14 out of 21) overall. Mem-LZerD thus overall outperformed the 33.3% success rate of JabberDock on preoriented inputs, despite requiring no expensive molecular dynamics simulation, while still performing well on targets with only predicted orientations.

### Docking results on the new transmembrane protein dataset

On the additional transmembrane benchmark unbound test set (Table 2), which included both α-helical and non-α-helical transmembrane proteins such as β-barrels, 35 of 44 targets (79.5%) were modeled by Mem-LZerD to at least acceptable CAPRI quality. This is broadly consistent with the performance demonstrated by Mem-LZerD on the smaller Memdock benchmark, which only included pairs of α-helical transmembane protein chains with an interface immersed in the membrane. Although this new benchmark includes full complexes rather than just pairs of chains and thus might be expected to offer increased opportunities to establish surface complementarity at the protein-protein interface, unconstrained rigid-body docking is not generally able to model these targets. Regular LZerD was only able to model 6 out of 44 (13.6%) targets in this benchmark to at least acceptable CAPRI quality. As indicated by the resulting low or zero 𝑓_nat_ values shown in the Regular LZerD Unbound column group of Table 2, the membrane environment severely interferes with recognition of the protein-protein interface. For example, in the case of the zinc metalloprotease mjS2P (PDB 3B4R), discussed in detail in Case Study 1, regular docking missed the interface entirely, while Mem-LZerD yielded a model of acceptable CAPRI quality. Mem-LZerD is of course dependent on the accuracy of its predicted subunit orientations, and this is reflected in the result for the bacterial magnesium transporter MgtE (PDB: 2YVX), where unbound docking succeeded but bound docking did not. In the absence of the full complex structure, PPM did not orient the subunits within the tilt angle cutoff used by Mem-LZerD. Thus, while acceptable poses were sampled in the full LZerD search, they were not emitted by Mem-LZerD as they all violated the tilt angle constraint. On the other hand, PPM oriented the unbound subunit structures appropriately. PPM, while a capable prediction method, is not explicitly designed to yield bound-state orientations from unbound-state structures. Thus, future developments of membrane docking should include an orientation prediction method designed for that purpose.

### Docking results on the peripheral membrane protein dataset

On the peripheral benchmark unbound test set (Supplementary Table S1), 15 of 92 targets (16.3%) were modeled to at least acceptable CAPRI quality within the top 10 models. This success rate is substantially lower than what was observed for transmembrane docking (61.9% in Table 1), as might be expected from the intuition that peripheral membrane proteins should be more difficult to orient to the membrane, especially when the subunits are oriented individually. Indeed, an unconstrained docking search sampled poses of at least acceptable CAPRI quality for 75 of 92 peripheral targets (81.5%) out of around 3 million total decoys typically for each target, as shown in the LZerD Sampling column of Supplementary Table S1. Among the sampled decoys generated, the standard LZerD pipeline was able to rank poses of at least acceptable CAPRI quality in the top 10 for 53 of 92 targets (57.6%), which is substantially higher than the 16.3% by Mem-LZerD, demonstrating that methods designed for soluble proteins are better at selecting good peripheral complex poses than the state-of-the-art PPM is at orienting the subunits. Figure 3 illustrates the particular challenges involved in docking peripheral membrane proteins. Figure 3a compares the modeling performance of regular LZerD to that of Mem-LZerD with predicted orientations, in terms of I-RMSD. For 59 out of 92 cases (64.1%), regular LZerD yielded a better (smaller) I-RMSD.

**Figure 3.**
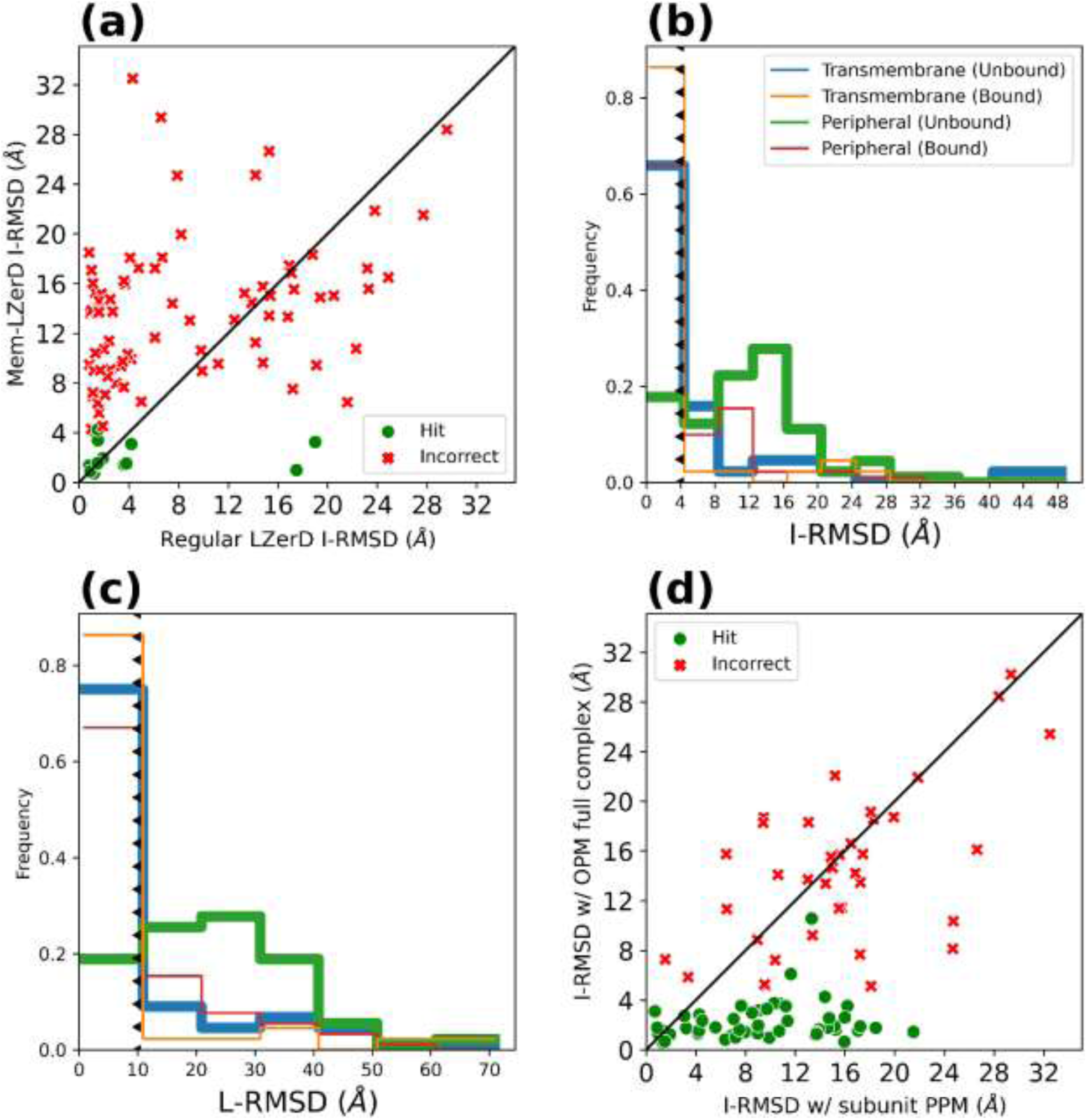
Comparison of peripheral membrane protein docking outcomes. **(a)** I-RMSD of Mem-LZerD using predicted orientations vs regular LZerD. Green: targets where Mem-LZerD was able to generate models of at least acceptable CAPRI quality in the top 10 when using predicted subunit orientations. Red: targets where Mem-LZerD failed to generate models of at least acceptable CAPRI quality in the top 10 when using predicted orientations. **(b)** I-RMSD best in top-10 distribution for transmembrane complexes vs peripheral membrane complexes. The triangle-dashed lines mark the CAPRI acceptable threshold for each metric, with the triangles pointing toward the acceptable side. **(c)** The same as (a), but for L-RMSD. **(d)** Performance of Mem-LZerD on peripheral membrane proteins using predicted versus ground truth membrane orientations. X-axis: the best I-RMSD among the top 10 models yielded by the Mem-LZerD pipeline when using the blind PPM subunit orientations. Y-axis: the best I-RMSD among the top 10 models yielded by the Mem-LZerD pipeline when using ground truth subunit orientations by superimposing to the oriented OPM complex. Green: targets where Mem-LZerD was able to generate models of at least acceptable CAPRI quality in the top 10 when using ground truth orientations. Red: targets where Mem-LZerD failed to generate models of at least acceptable CAPRI quality in the top 10 when using ground truth orientations.

Figures 3b and 3c show the Mem-LZerD output model quality for peripheral complexes as compared to transmembrane complexes in terms of the I-RMSD and L-RMSD CAPRI measures, respectively. While the distribution of each measure peaks within the CAPRI cutoffs of 4 Å and 10 Å, respectively, for transmembrane complexes cases, this is not the case for the peripheral complex targets. For the peripheral targets, the quality distributions peak far outside the corresponding acceptability thresholds. A comparison in terms of the remaining CAPRI measure, 𝑓_nat_, is shown in Supplementary Figure S1. To investigate what effect improvements to the subunit orientation predictions could have on the accuracy of the docking predictions, we recomputed the peripheral membrane protein targets using their bound orientations by superimposing the unbound subunits into the native structure taken from the OPM database, in the same way as in the transmembrane comparison (Figure 3d). I-RMSD improved when using bound (ground truth) orientations from OPM for 69 out of 92 cases (75.0%). With the bound orientations, models of at least acceptable CAPRI quality were built for 59.3% of cases within the top 10 (green dots in Figure 3d), while only 16.3% yielded acceptable models within the top 10 when using predicted subunit orientations by PPM. Since peripheral membrane proteins are by definition not immersed in the membrane, they are physically closer to soluble proteins than transmembrane proteins. This disparity confirmed that most of the peripheral targets are not sufficiently well oriented by PPM to dock using this method of constraining the search space. Supplementary Table S1 reflects this comparison in the Unbound Blind Mem-LZerD and Unbound Preoriented Mem-LZerD column groups. Future expansions of PPM or the inclusion of additional information from experiments which provide accurate orientation information, have the potential to change the outcome of peripheral complex docking from likely unsuccessful to likely successful.

### Case Study 1: Zinc metalloprotease mjS2P (PDB 3B4R)

Highlighted in Figure 4a is the zinc metalloprotease mjS2P complex from *Methanocaldococcus jannaschii*, a protease capable of cleaving proteins in the lipid membrane environment [49]. This complex is a key part of signaling by regulated intramembrane proteolysis, a signaling mechanism which has been established as conserved from bacteria to humans [50]. While this target was not tested in the Memdock or JabberDock studies, a run of AlphaFold as a substitute failed to model this complex, generating a model of incorrect CAPRI quality with an I-RMSD of 19.1 Å, and L-RMSD of 43.2 Å, and an 𝑓_nat_ of 0.01, shown in the right of Figure 4a. Although this model looks passably folded, it does not appear to even contain an appropriate transmembrane axis for the full complex. Indeed, the proper transmembrane axes of the two subunits are roughly perpendicular in this model, rather than roughly parallel. Mem-LZerD on the other hand was able to generate a model of acceptable CAPRI quality with an I-RMSD of 2.3 Å, and L-RMSD of 5.4 Å, and an 𝑓_nat_ of 0.49, shown in the left of Figure 4a. While the orange deviation lines show that the regions away from the protein-protein interface are not as precisely modeled, the interface residues are properly located. The backbone deviations away from the interface are largely parallel to the plane of the membrane and do not entail substantial dis-immersion. The inclusion by Mem-LZerD of reversed orientations was key to the successful modeling of this target, as otherwise Mem-LZerD would have only yielded incorrect models.

**Figure 4.**
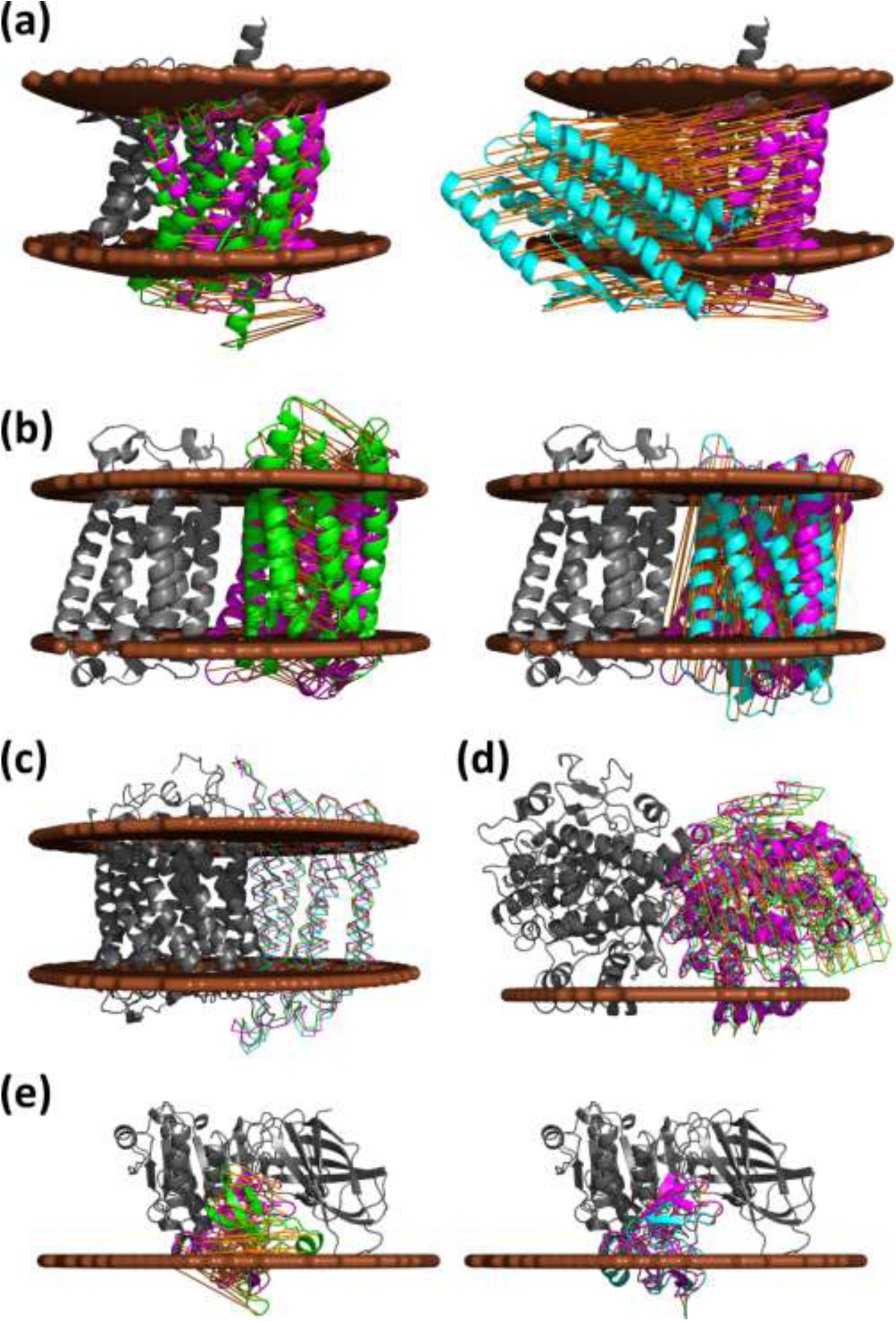
Example modeling of membrane protein complexes. Gray: the native receptor structure. Magenta: the native ligand binding pose. Brown: the membrane locations calculated by PPM. Orange: Cα deviations of one subunit when the other is superimposed. **(a)** Zinc metalloprotease mjS2P (PDB: 3B4R). Left: Mem-LZerD yielded a model (green) with an I-RMSD of 2.3 Å, and L-RMSD of 5.4 Å, and an 𝑓_nat_ of 0.49. Right: AlphaFold yielded a model (cyan) with an I-RMSD of 19.1 Å, and L-RMSD of 43.2 Å, and an 𝑓_nat_ of 0.01. **(b)** Elongation of very long chain fatty acids protein 7 (PDB: 6Y7F). Left: Mem-LZerD yielded a model (green) with an I-RMSD of 3.3 Å, and L-RMSD of 14.0 Å, and an 𝑓_nat_ of 0.38. Right: AlphaFold yielded a model (cyan) with an I-RMSD of 20.2 Å, and L-RMSD of 35.0 Å, and an 𝑓_nat_ of 0.00. **(c)** Poses of medium CAPRI quality generated by Mem-LZerD for cytochrome-c oxidase (PDB: 1M56). Cyan: first hit, an I-RMSD of 1.5 Å, and L-RMSD of 2.4 Å, and an 𝑓_nat_ of 0.71. Green: second hit, with an I-RMSD of 1.2 Å, and L-RMSD of 1.9 Å, and an 𝑓_nat_ of 0.84. Orange: displacements between the two poses, with an RMSD of 2.5 Å RMSD and maximum of 3.6 Å. **(d)** Prostaglandin G/H synthase 2 (PDB: 1CX2). Mem-LZerD yielded a model of acceptable CAPRI quality (green) with an I-RMSD of 3.2 Å, and L-RMSD of 6.4 Å, and an 𝑓_nat_ of 0.26, while doping with ground truth subunit orientations yielded a model of medium CAPRI quality (cyan) with an I-RMSD of 1.7 Å, an L-RMSD of 2.7 Å, and an 𝑓_nat_ of 0.51. Blind and doped model displacements are shown in orange and red respectively. **(e)** Pancreatic triacylglycerol lipase/colipase complex (PDB 1LPA). Left: Mem-LZerD yielded an incorrect model with an I-RMSD of 10.7 Å, and L-RMSD of 18.4 Å, and an 𝑓_nat_ of 0.02. Right: Mem-LZerD doped with ground truth subunit orientations yielded a model of acceptable CAPRI quality (cyan) with an I-RMSD of 3.7 Å, an L-RMSD of 5.0 Å, and an 𝑓_nat_ of 0.54.

### Case study 2: Elongation of very long chain fatty acids protein 7 (PDB 6Y7F)

The next case study, shown in Figure 4b, is the human ELOVL7 elongase complex, which is responsible for catalyzing the fatty acid elongation cycle [52] and is involved in a variety of serious genetic diseases in humans [53–55]. In the ELOVL7 homodimer observed by experiment, the substrate binds deep in a narrow tunnel formed by the structure of one subunit. The substrate binding site is not directly formed by the protein-protein interaction interface. The active site and interaction surface of this enzyme were noted to not be well-conserved in 2020 when its PDB entry 6Y7F was deposited, and modern template search techniques such as HHpred [51], using database PDB_mmCIF70_17_Apr, found no hits other than PDB 6Y7F itself. While this target was not tested in the Memdock or JabberDock studies, a run of AlphaFold as a substitute failed to model this complex, generating a model of incorrect CAPRI quality with an I-RMSD of 20.2 Å, and L-RMSD of 35.0 Å, and an 𝑓_nat_ of 0.00, shown in the right of Figure 4b. While the experimentalists observed the complex in a head-to-tail dimer both in crystal and in solution, AlphaFold instead generated head-to-head dimer models; in other words, AlphaFold placed one of the subunits upside down relative to the native complex structure, as indicated by the orange deviation lines. Mem-LZerD on the other hand was able to generate a model of acceptable CAPRI quality with an I-RMSD of 3.3 Å, and L-RMSD of 14.0 Å, and an 𝑓_nat_ of 0.38, shown in the left of Figure 4b. The inclusion by Mem-LZerD of reversed orientations was key to the successful modeling of this target as well, as otherwise Mem-LZerD would have only yielded incorrect models.

### Case study 3: Cytochrome-c oxidase (PDB 1M56)

The next example (Figure 4c) is a cytochrome-c oxidase complex from *Cereibacter sphaeroides*, a transmembrane enzyme found in the inner mitochondrial and cell membranes of bacteria [56]. For this target, both Memdock and JabberDock failed to retrieve a model of acceptable CAPRI quality in unbound docking, yielding only models of incorrect CAPRI quality within the top 10 models. Mem-LZerD on the other hand was able to generate a two models of medium CAPRI quality, one with an I-RMSD of 1.5 Å, and L-RMSD of 2.4 Å, and an 𝑓_nat_ of 0.71, and another with an I-RMSD of 1.2 Å, and L-RMSD of 1.9 Å, and an 𝑓_nat_ of 0.84. These two models thus both exceeded the criteria for acceptability. Mem-LZerD poses such as these can, for example, be used to instantiate molecular dynamics simulations with multiple reasonable initial conditions.

### Case study 4: Prostaglandin G/H synthase 2 (PDB 1CX2)

The last two examples are from the peripheral membrane protein dataset. Figure 4d shows a murine prostaglandin G/H synthase 2 (PTGS2) homodimer complex, a peripheral membrane enzyme in the nuclear and endoplasmic reticulum membranes of mice [57]. PTGS2 is involved in inflammatory response pathways, and interactions with homologous human proteins are the molecular basis of many non-steroidal anti-inflammatory drugs (NSAIDs) [58, 59]. PTGS2 binds to membranes peripherally via a helical amphipathic domain [60], which the protein-protein interface of PTGS2 does not include [61]. For this target, Mem-LZerD using predicted orientations from PPM was able to produce a model of acceptable CAPRI quality with an I-RMSD of 3.2 Å, and L-RMSD of 6.4 Å, and an 𝑓_nat_ of 0.26. With the correct orientation information taken from the full-complex orientation from OPM, Mem-LZerD yielded a model of medium CAPRI quality with an I-RMSD of 1.7 Å, an L-RMSD of 2.7 Å, and an 𝑓_nat_ of 0.51.

### Case study 5: Pancreatic triacylglycerol lipase with colipase (PDB 1LPA)

The last case study (Figure 4e) is a human pancreatic triacylglycerol lipase (PTL), a peripheral membrane protein involved in the metabolism of fats [62, 63], complexed with a porcine colipase (CLPS), shown in magenta in Figure 4e, a cofactor which facilitates membrane anchoring [64]. . Due to the presence of amphipathic bile salts in the physiological environment of PTL, the smaller CLPS is required as a cofactor to facilitate a binding mode with the catalytic site of PTL in sufficiently close contact with the membrane [64] to facilitate its function [62, 63]. For this target, Mem-LZerD using inputs blindly oriented by PPM was unable to produce a model of acceptable CAPRI quality, with the best of the top-10 models only reaching an I-RMSD of 10.7 Å, and L-RMSD of 18.4 Å, and an 𝑓_nat_ of 0.02, as shown in the left of Figure 4e. In this incorrect model, CLPS was placed at essentially the correct binding site, but is rotated about an axis roughly parallel to the membrane, completely disrupting the modeling of the native protein-protein interface. The blind orientation predictions of the subunits by PPM did not properly characterize the feasible region of docking poses for this target. When regular LZerD without constraints was used, we found that a correct binding pose with an I-RMSD of 3.7 Å was sampled, but it was not selected by the scoring function within top-10 models, instead yielding an even worse an I-RMSD of 22.3 Å, an L-RMSD of 51.8 Å, and an 𝑓_nat_ of 0.00 within the top 10. When instead modeling using the full-complex orientation from OPM, Mem-LzerD yielded a model of medium CAPRI quality with an I-RMSD of 3.7 Å, an L-RMSD of 5.0 Å, and an 𝑓_nat_ of 0.54, as shown in the right of Figure 4e. Thus, in this case, the correct orientation was required for a correct pose to be discovered by the scoring function.

### Preparation of molecular dynamics with membrane docking in the LZerD webserver

Due to the limited ability of models like AlphaFold to sample many interaction poses out-of-the-box, biologists interested in modeling the dynamics of proteins in a membrane may need to arrange individual protein chains to begin modeling the states desired, or even reasonable states. This difficulty is exemplified by Case Study 2 above, where despite experimental evidence to the contrary, the AlphaFold models of ELOVL7 took on entirely opposite orientations. Mem-LZerD can assemble ELOVL7 into reasonable bound poses which can then be modeled in explicit lipid solvent and simulated. Mem-LZerD is freely accessible via the LZerD webserver for protein docking, available at https://lzerd.kiharalab.org [16, 17], which allows users without high-performance computers or without computing experience to easily model the assembly of protein complexes. Figure 5a shows the input preparation for modeling poses of ELOVL7 using the LZerD webserver. Input files for docking can be taken directly from the OPM website https://opm.phar.umich.edu/) [44] or can be oriented using the PPM website or locally-installed application. The webserver allows users to specify residue-based distance constraints on the docking, and more information on this and all functionality can be found in previous works [16, 17] and the About pages on the webserver. However, in this case the default options are sufficient, and we can proceed to submit the docking. Shown in Figure 5b is the results page of this docking. Note that the ligand pose centroids, shown as spheres, highlight the focus of the membrane docking search. From here, we can download our desired models and upload them to CHARMM-GUI (https://www.charmm-gui.org) [46]. Figure 5c highlights the steps of processing a docked model using CHARMM-GUI. In this instance, we selected PLPC lipids and the NAMD output format. Figure 5d shows ELOVL7 solvated in the periodic membrane, and Supplementary Video S2 shows the equilibration and a 1ns production molecular dynamics trajectory of this system. With these together, we see that the dynamics simulation proceeds smoothly.

**Figure 5.**
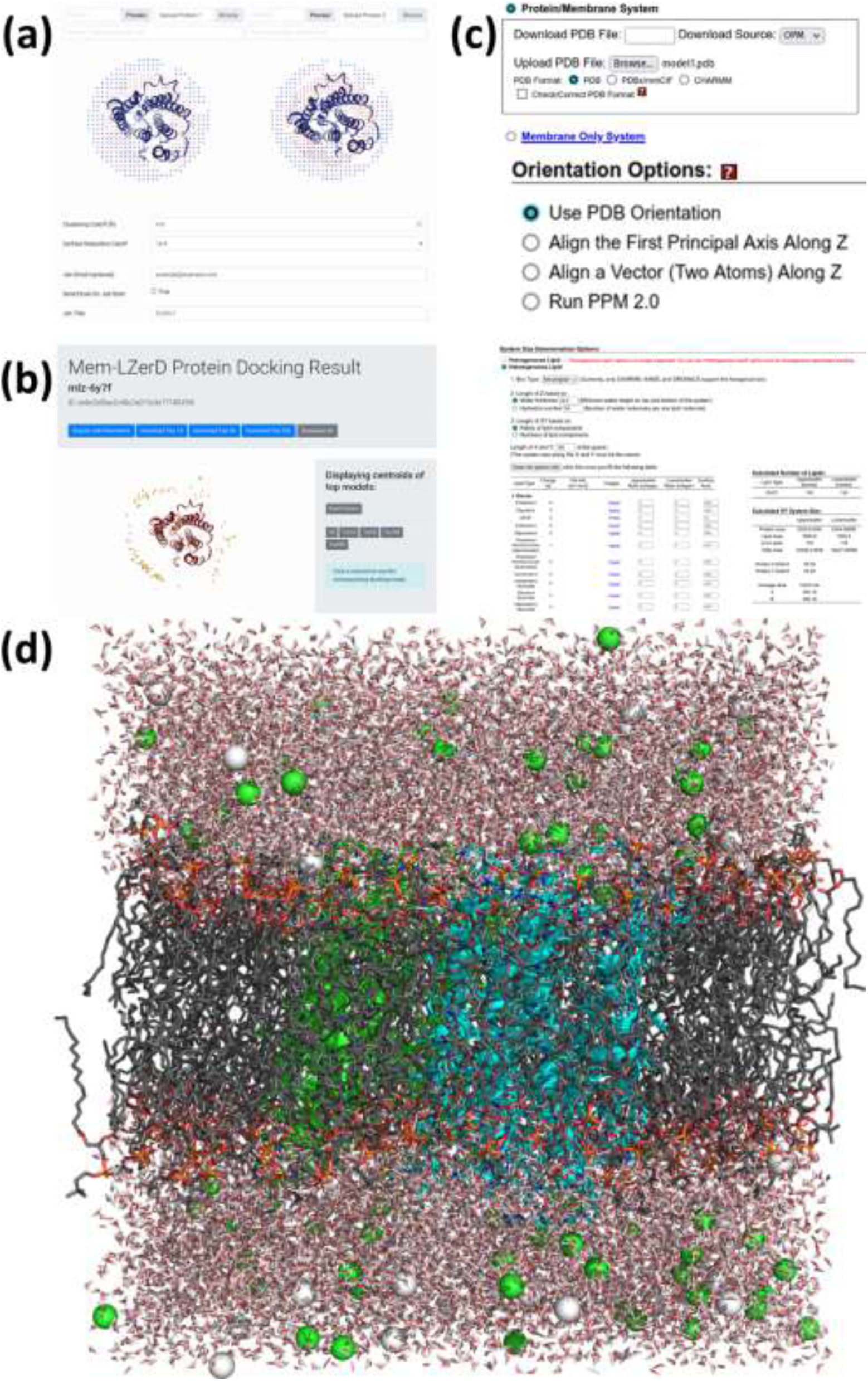
Example workflow using the LZerD webserver to prepare and run a molecular dynamics simulation of a docked model of target elongation of very long chain fatty acids protein 7 (PDB 6Y7F) in the membrane. **(a)** PPM-oriented receptor and ligand protein models uploaded to the LZerD webserver submission page. Default settings are used in the absence of residue distance information. **(b)** Mem-LZerD docking results shown in the LZerD webserver results viewer. Each sphere shows the centroid of a sampled pose. **(c)** Certain options used in generating NAMD input files using CHARMM-GUI. A PPM-oriented PDB file was uploaded, and its precomputed orientation was kept. PLPC lipids and a box size of 100 were used to construct the explicit simulation-ready files. **(d)** The final prepared coordinates for simulation.

## MATERIALS AND METHODS

### Dataset construction

The transmembrane dataset used in this work was constructed by starting from 4218 nominally transmembrane protein structures from the OPM database [45]. Monomers were excluded, and entries were clustered at 25% sequence identity using MMseqs2 [65], yielding 390 complexes. Structure-based clustering with an MM-align [66] TM-score cutoff of 0.50 yielded 85 complexes. From these, complexes which were not oriented inside the membrane and for which the publication did not indicate that the complex was in fact studied in a membrane were excluded, or which were otherwise beyond the scope of Mem-LZerD, yielding the final transmembrane dataset of 64 complexes. 20 complexes from this dataset were separated to serve as the training set for the scoring function, while the remaining 44 complexes served as the test set. For all targets, AlphaFold v2.2.2 [12] with v2 weights and without template search was used to generate unbound subunit models.

The peripheral membrane complex dataset was constructed in a similar manner to the transmembrane dataset. Starting from 1607 complexes listed by OPM as “monotopic/peripheral”, 147 complexes with two subunits and each interacting with the membrane were extracted. After clustering as with the transmembrane dataset, 92 complexes remained, constituting the final peripheral membrane complex dataset. Peripheral complexes were not used to train the scoring function, and thus this dataset was not partitioned.

### Rigid-body docking of subunits with regular LZerD

Each ligand protein structure subunit was docked with the receptor subunit using LZerD [14, 67]. LZerD is a shape complementarity-based rigid-body docking algorithm which employs a soft representation of the molecular surface to tolerate some small differences between the bound and unbound conformations. Using geometric hashing-based surface searching, LZerD generates many candidate docking poses, which are then scored according to the shape complementarity of their interfaces. These initial candidate poses, before any truncation or clustering, are referred to as the poses sampled by LZerD. For a given pair of input subunits, the set of docked models generated by LZerD was ordered by the LZerD shape score and truncated to a set of 50,000 models. This set was then clustered with an RMSD cutoff of 4.0 Å. The final stage of the regular LZerD pipeline is a rescoring according to the ranksum scoring function [15, 74, 75].

### Membrane-aware pose constraints

The orientations in the membrane of models to be docked were predicted using the Positioning of Proteins in Membranes (PPM) software package [44, 45] using default settings. PPM outputs an oriented version of each input structure and also predicts the interior and exterior membrane surfaces. The separately oriented subunits were assumed to share a membrane midplane. These predicted orientations are then used to augment the LZerD docking search. In LZerD, sample points on the protein surface are matched to each other in the geometric hashing discussed above. To accomplish this, reference frames are constructed using pairs of nearby points on the surface in combination with their surface normals. The other surface points in a fixed neighborhood of the origin of this reference frame are translated and rotated such that their basis is changed to the reference frame, resulting in a fingerprint of the molecular surface. This procedure was implemented using a space-partitioning tree data structure [68], which is designed to efficiently search for points in Euclidean space by querying only relevant regions of space. By selecting appropriate planes with which to divide the space of hashed points, individual queries in the accumulation and recognition stage of the geometric hashing are made in sublinear time.

In Mem-LZerD, this data structure is augmented with the distance of each point to the membrane midplane. These height values for each point come directly from the output of PPM and are fixed once the geometric hashing data structure has been built at the beginning of the LZerD search. During a query to the geometric hashing data structure, hashed points tagged with a height different from the query height by more than the cutoff will not be emitted, and thus not contribute to the match vote accumulation of the geometric hashing. Unlike the 3-dimensional coordinates of the sample points, the height value is not transformed during the change of basis, and the internal structure thus differs from otherwise similar lifting transform techniques that bridge related entities in computational geometry [69]. This augmentation facilitates efficient imposition of constraints in terms of the relative midplane distance, essentially pruning the search space analyzed. For Mem-LZerD we used a relative midplane distance of 8 Å. As surface matches satisfying the relative midplane distance constraint are found, their docked orientations relative to their predicted orientations were further checked to satisfy a relative midplane axis constraint of 0.4 radians. The base orientation axes are fixed at the start of docking based on PPM, and this tilt angle check does not use the properties of the geometric hashing data structure, as it requires that the full rigid-body transformation have already been calculated. While these are the same numerical thresholds as were previously used by Memdock, we note that they are comparable but different quantities than those constrained in Memdock. We show a validation of these hyperparameters on the training set in Figure 2c based on the recall of models of at least acceptable CAPRI quality and the enrichment factor (EF, the ratio of the fraction of hits in the decoy set after constraining to the fraction of hits in the decoy set before constraining) of the pose space constrained to. When the parameter combinations were ordered by their corresponding EF values, we found that the recall of the constrained decoy set declined dramatically. As illustrated in Figure 2ab, nearly all of the search space outside this feasible region consists of incorrect models. Additionally, the implementation of the height constraint into the geometric hashing results in a running time 74 times faster compared to regular LZerD.

### Docked model scoring

For selecting docked poses, we constructed a combined scoring function derived from methods used in the scoring of IDP-LZerD and Flex-LZerD [34–36]. This scoring function uses a bagged decision tree estimator [70] to combine the knowledge-based scoring functions GOAP [71], DFIRE [72], and ITScorePro [73], which are the three re-scoring functions combined in the usual ranksum score for competitive LZerD docking [15, 74, 75], as well as LZerD initial docking score, the cluster size from the 4 Å RMSD clustering normally used with LZerD, membrane transfer energy [76], and order statistic terms (𝑂𝑆_(1)_, 𝑂𝑆_(2)_, and 𝑂𝑆_(3)_) highlighting the most-outlying values among the other scoring terms for each model. The LZerD shape complementarity scores and the cluster sizes are part of the regular LZerD output.

The order statics terms allow models with any outlying component scores a better chance at inclusion among the top-ranked models, even if some other component scores are low or generally disagree with each other. Each model score for all docked poses of a target is first standardized in the usual way into a Z-score. For example, the mean 𝜇 and standard deviation 𝜎 of all ITScorePro scores 𝑥_𝑖_ of models of sucrose-specific porin are calculated, and the Z-score of model 𝑖 will be 𝑧_𝑖_ = (𝑥_𝑖_ − 𝜇)/𝜎. For scoring terms designed such that more positive values are more favorable, e.g. the LZerD shape complementarity term and cluster sizes, the values were multiplied by −1 so that lower values would uniformly be more favorable across all component scores. Thus, for all the final standardized scoring terms, a more negative Z-score is more favorable. From these, the order statistic terms were calculated for the Z-scores of the scoring terms. To compute e.g. the order statistic *OS(1)* of a model, we first check the Z-sores of all the scoring terms for the model and select the smallest (the most favorable) Z-score. Following the same logic, *OS(2)* and *OS(3)* are the second and the third smallest Z-scores among all the Z-scores of all the scoring terms. A first-order version of this approach was previously used successfully in IDP-LZerD for disordered proteins [36], and a third-order version was used successfully in Flex-LZerD for flexible proteins [34, 35]. Here, we use the third-order version. For example, if a model has Z-scores of −1.7, −2.4, and −0.5 for the shape score, GOAP, and ITScorePro, respectively, and some arbitrary positive-valued Z-scores for the component scores, then *OS(1), OS(2)*, and *OS(3)* for that model will be −2.4, −1.7, and −0.5, respectively.

The component scores GOAP, DFIRE, ITScorePro, ranksum, the LZerD shape score, LZerD cluster size, membrane transfer energy, 𝑂𝑆_(1)_, 𝑂𝑆_(2)_, and 𝑂𝑆_(3)_, were then finally combined in the bagged estimator. Bagging is an ensemble method which constructs multiple estimators using subsets of the training data, and then averages their individual outputs to produce a more robust classifier [70]. To optimize the bagged ensemble, docked models were generated for each target in the training dataset using the same procedure as above. CAPRI statistics were then calculated for each model. Each model was labeled positive if it was of acceptable CAPRI quality, and negative if it was not. Each model was then weighted to balance the number of positive and negative cases within each target. Then, the bagged estimator was trained using all the models for all the training targets. The resulting scoring function, with ranksum component scores used to break ties, was used to rescore the docked models after the clustering stage.

## Supporting information

supplemental-materials-misc

## ACKNOWLEDGEMENTS

The authors thank Daipayan Sarkar for his contribution to an earlier form of the project. The authors are grateful to Information Technology at Purdue, West Lafayette, Indiana for providing computational resources. This publication was made possible with computational resource support from the Purdue Institute of Inflammation, Immunology and Infectious Disease (PI4D).

## DECLARATIONS OF INTEREST

The authors declare no conflict of interest.

[TAB]

