## supplemental-materials-misc for "Assembly of Protein Complexes In and On the Membrane with Predicted Spatial Arrangement Constraints"

**Supplemental materials for**
### Assembly of Protein Complexes In and On the Membrane with Predicted Spatial Arrangement Constraints

Charles Christoffer[1] and Daisuke Kihara[1,2,*]

[1] Department of Computer Science, Purdue University, West Lafayette, Indiana, 47907, USA
[2] Department of Biological Sciences, Purdue University, West Lafayette, Indiana, 47907, USA

* Corresponding author.


| Target | Unbound Blind Mem-LZerD | | | Unbound Preoriented Mem-LZerD | | | Regular LZerD | | |
|---|---|---|---|---|---|---|---|---|---|
| | I-RMSD (Å) | L-RMSD (Å) | $f_{nat}$ | I-RMSD (Å) | L-RMSD (Å) | $f_{nat}$ | I-RMSD (Å) | L-RMSD (Å) | $f_{nat}$ |
| 1bwn | (18.3) | (46.2) | (0.00) | (18.5) | (34.2) | (0.00) | (18.8)/1.6 | (54.4)/4.2 | (0.00)/0.54 |
| 1cx2 | 3.2 | 6.4 | 0.26 | 1.7 | 2.7 | 0.51 | (19.0)/1.7 | (61.9)2.7 | (0.00)/0.67 |
| 1djt | 3.1 | 6.7 | 0.41 | 2.7 | 4.7 | 0.65 | (4.2)/0.9 | (11.1)0.9 | (0.41)/0.68 |
| 1doa | (15.3) | (23.7) | (0.00) | 1.6 | 3.0 | 0.65 | 1.3/1.3 | 1.7/1.7 | 0.74/0.74 |
| 1dvp | (14.5) | (36.5) | (0.00) | (13.4) | (24.8) | (0.06) | (13.9)/9.8 | (30.7)29.0 | (0.06)/0.06 |
| 1dyn | (9.6) | (22.5) | (0.23) | (5.2) | (10.1) | (0.23) | (14.8)/1.7 | (42.1)/3.9 | (0.00)/0.65 |
| 1e6c | (6.5) | (11.8) | (0.04) | (11.3) | (21.5) | (0.01) | 5.0/5.0 | 9.2/9.2 | 0.14/0.14 |
| 1eci | 1.4 | 3.0 | 0.80 | 1.3 | 2.4 | 0.70 | 1.41.4 | 3.0/2.5 | 0.80/0.80 |
| 1efv | (9.9) | (16.6) | (0.01) | 1.0 | 1.7 | 0.79 | 4.2/1.3 | 9.2/2.6 | 0.19/0.57 |
| 1fle | 0.7 | 1.5 | 0.75 | 3.1 | 7.9 | 0.28 | 1.2/0.7 | 3.1/1.7 | 0.77/0.82 |
| 1g5z | (13.6) | (22.6) | (0.05) | 1.2 | 2.0 | 0.78 | 0.9/0.9 | 1.2/1.2 | 0.86/0.86 |
| 1gxd | (9.3) | (22.5) | (0.01) | 3.2 | 3.9 | 0.41 | 3.4/3.2 | 3.6/7.8 | 0.33/0.33 |
| 1joc | (15.2) | (33.2) | (0.00) | 1.9 | 3.4 | 0.75 | 1.8/1.8 | 2.3/2.3 | 0.79/0.80 |
| 1lpa | (10.7) | (18.4) | (0.02) | 3.7 | 5.0 | 0.54 | (22.3)/3.7 | (51.8)/4.8 | (0.00)/0.37 |
| 1lsh | (17.4) | (29.7) | (0.00) | (15.8) | (26.4) | (0.00) | (16.9)/4.5 | (34.2)/7.9 | (0.00)/0.13 |
| 1m7r | (10.6) | (27.7) | (0.00) | (14.1) | (57.8) | (0.00) | (9.8)/1.3 | (38.1)/4.9 | (0.00)/0.71 |
| 1moz | (26.6) | (65.4) | (0.00) | (16.1) | (28.5) | (0.29) | (15.3)/14.7 | (21.2)/30.8 | (0.00)/0.00 |
| 1ocu | (28.4) | (39.2) | (0.00) | (28.4) | (37.2) | (0.00) | (29.6)/26.4 | (46.7)/27.1 | (0.00)/0.00 |
| 1of8 | (16.0) | (28.0) | (0.00) | 0.7 | 1.4 | 0.87 | 3.7/2.7 | 11.1/6.5 | 0.30/0.32 |
| 1qge | 1.4 | 1.8 | 0.75 | 1.5 | 2.2 | 0.65 | 1.4/1.4 | 1.8/1.8 | 0.75/0.75 |
| 1tu5 | (15.2) | (31.2) | (0.00) | (22.1) | (42.3) | (0.00) | (13.3)/10.8 | (27.6)/24.0 | (0.00)/0.01 |
| 1tx2 | 1.3 | 4.2 | 0.62 | 0.6 | 2.0 | 0.90 | 0.8/0.8 | 3.2/2.3 | 0.80/0.92 |
| 1u5d | (17.2) | (40.8) | (0.00) | (13.5) | (30.5) | (0.00) | 4.8/4.2 | 9.8/12.3 | 0.28/0.42 |
| 1u7n | (14.6) | (36.4) | (0.01) | 1.8 | 3.3 | 0.62 | 1.6/1.6 | 2.1/4.3 | 0.51/0.65 |
| 1v02 | (9.5) | (27.1) | (0.00) | (18.7) | (38.7) | (0.00) | 0.8/0.8 | 3.5/3.5 | 0.85/0.85 |
| 1ve9 | 1.4 | 3.6 | 0.67 | 0.9 | 2.7 | 0.90 | 3.6/1.0 | 9.1/2.2 | 0.22/0.64 |
| 1vf6 | 4.2 | 7.6 | 0.21 | 1.3 | 2.5 | 0.79 | 1.1/1.1 | 1.9/1.9 | 0.62/0.62 |
| 1via | (5.6) | (10.8) | (0.20) | 1.8 | 4.3 | 0.80 | 1.6/1.5 | 2.5/2.0 | 0.82/0.80 |

| | | | | | | | | | |
|---|---|---|---|---|---|---|---|---|---|
| 1wdz | 3.4 | 7.4 | 0.23 | (5.8) | (13.2) | (0.03) | 1.5/1.5 | 4.0/4.0 | 0.51/0.51 |
| 1z9h | (17.2) | (26.8) | (0.00) | (7.7) | (23.5) | (0.07) | (23.2)/1.6 | (51.3)/3.7 | (0.00)/0.42 |
| 1zww | (4.3) | (12.0) | (0.13) | 2.8 | 5.4 | 0.18 | 1.0/1.0 | 2.9/2.9 | 0.86/0.86 |
| 2b3r | (8.0) | (19.4) | (0.21) | 1.4 | 3.9 | 0.83 | 3.0/0.9 | 10.0/1.8 | 0.58/0.75 |
| 2bng | 1.0 | 3.0 | 0.75 | 1.3 | 5.0 | 0.62 | (17.5)/1.0 | (45.9)/3.0 | (0.12)/0.75 |
| 2d2r | (11.4) | (25.0) | (0.05) | 2.3 | 2.6 | 0.68 | 2.4/2.2 | 2.8/2.8 | 0.64/0.58 |
| 2dfk | (11.6) | (20.1) | (0.01) | 6.1 | 6.3 | 0.68 | 6.1/6.1 | 8.6/8.6 | 0.73/0.73 |
| 2efl | (14.7) | (27.9) | (0.04) | 2.6 | 6.2 | 0.26 | 2.5/2.3 | 5.8/5.8 | 0.35/0.26 |
| 2fic | (18.5) | (47.5) | (0.02) | 1.8 | 3.9 | 0.36 | 0.8/0.8 | 3.2/3.2 | 0.89/0.89 |
| 2gkn | (9.0) | (28.0) | (0.07) | (8.9) | (30.8) | (0.00) | (9.9)/1.1 | (30.5)/5.4 | (0.00)/0.93 |
| 2hd0 | (16.8) | (26.4) | (0.00) | (14.2) | (23.8) | (0.04) | (17.1)/9.2 | (38.6)/13.6 | (0.00)/0.00 |
| 2hsj | (6.9) | (14.2) | (0.27) | 1.1 | 2.4 | 0.70 | 1.1/1.1 | 2.3/2.3 | 0.77/0.77 |
| 2q63 | (10.3) | (16.4) | (0.03) | 3.8 | 3.7 | 0.50 | 3.9/3.9 | 5.1/5.1 | 0.36/0.36 |
| 2qpt | (21.9) | (24.9) | (0.08) | (21.9) | (29.7) | (0.27) | (23.8)/19.2 | (31.5)/26.4 | (0.26)/0.00 |
| 2sqc | (15.8) | (48.7) | (0.07) | (11.4) | (30.6) | (0.02) | (14.8)/0.6 | (24.4)/1.7 | (0.00)/0.84 |
| 2v0o | (15.0) | (26.1) | (0.01) | (14.6) | (23.8) | (0.15) | (15.4)/12.7 | (24.6)/18.5 | (0.00)/0.09 |
| 2wuj | (6.4) | (11.2) | (0.13) | 0.8 | 1.2 | 0.87 | 1.5/1.3 | 3.1/2.3 | 0.59/0.50 |
| 2zsh | (9.0) | (17.8) | (0.05) | 1.3 | 2.1 | 0.76 | 1.2/1.1 | 2.0/1.8 | 0.65/0.64 |
| 3a36 | (11.2) | (26.1) | (0.00) | 3.5 | 11.0 | 0.46 | (14.2)/2.3 | (31.2)/5.5 | (0.00)/0.49 |
| 3ajm | (9.1) | (20.0) | (0.08) | 3.2 | 5.8 | 0.37 | 2.5/2.4 | 9.9/8.0 | 0.58/0.62 |
| 3caz | (10.7) | (17.6) | (0.01) | 1.5 | 2.5 | 0.80 | 2.0/2.0 | 4.4/4.4 | 0.54/0.54 |
| 3fgr | (13.8) | (25.7) | (0.01) | 1.2 | 2.4 | 0.67 | 1.0/1.0 | 1.9/1.9 | 0.83/0.83 |
| 3g7j | 0.9 | 1.7 | 0.69 | 1.8 | 2.9 | 0.53 | 0.9/0.9 | 1.7/1.7 | 0.69/0.69 |
| 3gny | 1.9 | 3.9 | 0.62 | 1.2 | 2.3 | 0.90 | 1.9/0.7 | 3.9/1.6 | 0.62/0.86 |
| 3hhm | (29.4) | (56.1) | (0.01) | (30.2) | (48.6) | (0.00) | (6.6)/3.3 | (13.1)/5.6 | (0.20)/0.38 |
| 3hl4 | 4.2 | 8.0 | 0.18 | 1.5 | 2.1 | 0.67 | 1.5/1.5 | 2.0/2.0 | 0.64/0.64 |
| 3icv | (13.9) | (36.4) | (0.04) | 1.7 | 3.2 | 0.74 | 1.6/1.6 | 2.1/2.1 | 0.55/0.55 |
| 3kl4 | (16.2) | (39.5) | (0.00) | 3.5 | 6.8 | 0.28 | 3.6/3.1 | 9.9/8.2 | 0.50/0.56 |
| 3lxr | (17.1) | (28.1) | (0.01) | 1.5 | 3.5 | 0.65 | 1.0/1.0 | 2.3/2.3 | 0.90/0.90 |
| 3mdk | (7.2) | (15.2) | (0.13) | 1.0 | 2.1 | 0.84 | 1.1/0.9 | 2.2/1.4 | 0.77/0.67 |
| 3nrd | 1.5 | 3.3 | 0.43 | 0.7 | 1.6 | 0.92 | 3.8/1.1 | 6.7/2.2 | 0.23/0.51 |

| | | | | | | | | | |
|---|---|---|---|---|---|---|---|---|---|
| 3ogh | 1.5 | 1.4 | 0.79 | (7.3) | (14.8) | (0.05) | 1.5/1.5 | 1.4/1.4 | 0.79/0.79 |
| 3rbb | (19.9) | (53.2) | (0.00) | (18.7) | (57.0) | (0.00) | 8.2/5.6 | 30.1/16.8 | 0.00/0.17 |
| 3w54 | (16.5) | (20.4) | (0.17) | (16.6) | (21.1) | (0.22) | (24.9)/16.2 | (48.3)/19.3 | (0.00)/0.02 |
| 3wxx | (13.3) | (20.5) | (0.05) | 10.6 | 8.0 | 0.44 | (16.8)/11.8 | (29.4)/15.5 | (0.01)/0.04 |
| 4bik | (18.1) | (43.2) | (0.00) | (5.1) | (13.7) | (0.09) | (6.7)/1.4 | (20.1)/2.8 | (0.00)/0.60 |
| 4bne | (8.5) | (16.8) | (0.01) | 3.0 | 7.1 | 0.36 | 2.3/2.3 | 7.3/6.8 | 0.82/0.61 |
| 4d9o | (9.4) | (20.0) | (0.03) | (18.3) | (45.3) | (0.03) | (19.1)/8.9 | (45.0)/12.4 | (0.00)/0.05 |
| 4h8s | (24.7) | (47.2) | (0.00) | (8.1) | (17.3) | (0.11) | (7.9)/6.0 | (17.7)/13.5 | (0.09)/0.09 |
| 4klr | (17.2) | (33.6) | (0.06) | 1.9 | 3.4 | 0.63 | (6.1)/1.5 | (12.8)/2.7 | (0.08)/0.69 |
| 4nsw | (32.5) | (66.5) | (0.01) | (25.4) | (47.9) | (0.00) | 4.3/4.3 | 9.6/9.7 | 0.32/0.32 |
| 4pus | (9.5) | (20.0) | (0.08) | (5.3) | (14.2) | (0.08) | (11.2)/1.1 | (35.6)/2.5 | (0.05)/0.55 |
| 4qn9 | (13.1) | (40.3) | (0.00) | (18.3) | (28.7) | (0.06) | (12.5)/2.5 | (33.1)/8.5 | (0.06)/0.44 |
| 4wpe | (13.7) | (35.0) | (0.09) | 1.4 | 2.7 | 0.54 | 1.6/1.6 | 4.1/4.1 | 0.34/0.34 |
| 4zhk | (21.5) | (37.2) | (0.00) | 1.4 | 1.6 | 0.73 | (27.7)/1.8 | (80.7)/4.1 | (0.00)/0.61 |
| 4zv5 | (9.7) | (19.1) | (0.04) | 3.3 | 3.0 | 0.64 | 3.5/3.2 | 3.7/2.9 | 0.79/0.64 |
| 5a52 | (13.4) | (37.0) | (0.00) | (9.2) | (26.0) | (0.00) | (15.3)/0.8 | (41.9)/2.2 | (0.00)/0.69 |
| 5c1f | (18.1) | (34.8) | (0.00) | (19.2) | (33.8) | (0.00) | 4.1/3.5 | 7.0/6.6 | 0.19/0.14 |
| 5dyy | (6.4) | (16.8) | (0.04) | (15.8) | (30.4) | (0.04) | (21.6)/2.2 | (79.4)6.7 | (0.00)/0.26 |
| 5fqu | (9.0) | (21.8) | (0.04) | 2.0 | 4.9 | 0.79 | 1.7/1.3 | 1.8/5.9 | 0.71/0.75 |
| 5jwa | (7.5) | (13.4) | (0.07) | 1.6 | 3.2 | 0.43 | (17.2)/1.4 | (36.3)/3.0 | (0.00)/0.62 |
| 5kk4 | (7.6) | (15.7) | (0.04) | 3.5 | 8.3 | 0.29 | 3.6/0.9 | 9.5/2.6 | 0.46/0.88 |
| 5oo7 | (15.0) | (32.0) | (0.00) | (15.2) | (27.2) | (0.00) | (20.5)/2.3 | (41.2)/4.4 | (0.00)/0.35 |
| 5w7b | (16.0) | (31.3) | (0.00) | 2.6 | 4.3 | 0.41 | 1.1/1.1 | 2.9/2.9 | 0.91/0.91 |
| 5w7c | (13.7) | (22.5) | (0.05) | 1.3 | 1.9 | 0.92 | 2.7/1.6 | 6.4/2.6 | 0.43/0.65 |
| 6b3i | (15.6) | (30.8) | (0.02) | (15.7) | (19.3) | (0.14) | (23.3)/13.4 | (42.8)/54.8 | (0.00)/0.02 |
| 6bym | (14.4) | (22.1) | (0.06) | 4.3 | 6.8 | 0.34 | (7.5)/3.3 | (17.4)/3.3 | (0.06)/0.44 |
| 6hln | (7.0) | (14.6) | (0.06) | 2.5 | 3.4 | 0.40 | 2.1/1.9 | 2.9/3.7 | 0.46/0.46 |
| 6i7s | (24.7) | (36.1) | (0.01) | (10.3) | (7.7) | (0.07) | (14.2)/9.4 | (39.0)/7.6 | (0.05)/0.04 |
| 6ikn | (10.4) | (24.3) | (0.08) | (7.2) | (11.9) | (0.22) | 1.3/1.3 | 2.3/2.3 | 0.73/0.73 |
| 6koi | (14.9) | (31.1) | (0.05) | (15.5) | (24.2) | (0.01) | (19.4)/9.4 | (43.1)/10.8 | (0.00)/0.00 |
| 6lcq | (4.5) | (10.6) | (0.26) | 2.3 | 4.5 | 0.72 | 1.9/1.5 | 3.4/1.9 | 0.74/0.74 |

| | | | | | | | | | |
|---|---|---|---|---|---|---|---|---|---|
| 6tlb | (15.5) | (24.1) | (0.00) | (11.4) | (14.8) | (0.04) | (17.3)/9.9 | (26.1)/13.4 | (0.00)/0.04 |
| 6vzd | (13.0) | (21.0) | (0.22) | (13.7) | (23.6) | (0.12) | (8.9)/6.5 | (13.8)/6.6 | (0.03)/0.06 |
| **Total successes:** | | | **15 (16.3%)** | | | **54 (58.7%)** | | | **53 (57.6%)** |

**Supplementary Table S1.** Docking performance of the 92 individual targets of the Flex-LZerD peripheral membrane complex benchmark set. The columns list the CAPRI measures for the best-I-RMSD model among the top 10 models by the scoring function for each docking method respectively. Parentheses indicate that no models of at least CAPRI-acceptable quality were ranked within the top 10. For the Regular LZerD column group, the CAPRI measures of the best sampled model by I-RMSD are given after the slash.

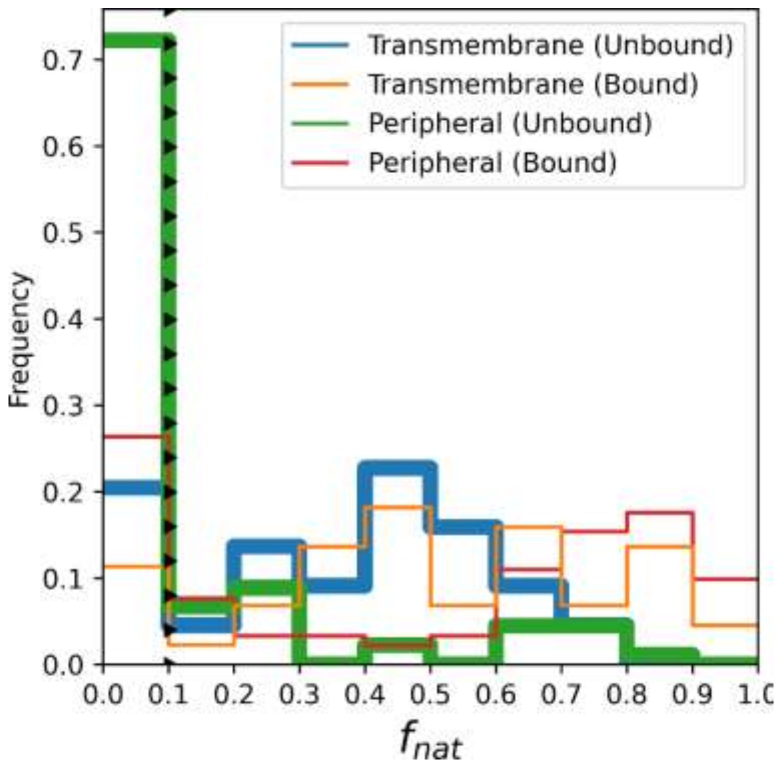


**Supplementary Figure S1.** $f_{\mathrm{nat}}$ best in top-10 distribution for transmembrane complexes vs peripheral membrane complexes. The triangle-dashed lines mark the CAPRI acceptable threshold of 0.10, with the triangles pointing toward the acceptable side.

**Supplementary Video S1.** Top 500 sampling of docking of two subunits of an intramembrane aspartate protease (PDB 4HYG, chains A and B). Sampled model centroids are shown as gold spheres. **Left, purple:** Sampling using regular LZerD. Many clearly impossible poses are sampled, including completely sideways orientations. **Right, cyan:** Sampling using Mem-LZerD. The poses sampled are more focused towards compatible orientations upright and in the membrane.

**Supplementary Video S2.** Example 1-nanosecond explicit-membrane molecular dynamics simulation instantiated from a Mem-LZerD model of the target elongation of very long chain fatty acids protein 7 (PDB 6Y7F) using a PLPC membrane instantiated by CHARMM-GUI.
